# Detection and Characterization of Magnetosome Chains In Magnetotactic Bacteria

**DOI:** 10.1101/2020.08.20.258723

**Authors:** Bharadhwaj Vinay, Bhavya Vijay, Surya K Raghunath

## Abstract

**Aim:** Magnetotactic bacteria are gram-negative, prokaryotic organisms which align themselves according to the Earth’s geomagnetic field. They contain organelles called magnetosomes which produce nano-magnetites by the mechanism of biomineralization. These nano-magnetites are arranged in the form of well-ordered chain called magnetosome chain, which are held together by intermolecular forces. The growth of these bacteria is stringent to specific oxygen gradient regions, where there is oxic-anoxic transition zone. In this paper, we report the cultivation and characterisation of magnetotactic bacteria that was accomplished under laboratory conditions and scanning electron microscope respectively.

**Methodology:** Bacterial soil samples were collected and cultivated under laboratory conditions using a precise and cost-effective media formulation using coffee bean extract and ferric chloride, which were to provide sufficient iron under the influence of external magnetic field. Experiments were conducted using media, deficient with ferric chloride and coffee bean solution, as control. Pour plate method was carried out for the growth of bacteria under the influence of external magnetic field provided on one-end of the petri-plate.

**Result:** We observed the growth of the cultivated bacteria in the proximity of the magnetic field only in the presence of iron in media. This emphasises that the media formulated was appropriate for the growth of bacteria under laboratory conditions. Scanning electron microscope analysis confirmed the presence of magnetosome chains in magnetotactic bacteria.

**Interpretation:** In this research, the bacteria were able to sustain in artificial oxygen-transition zones provided in the laboratory. The coffee bean solution contained quinic acid and succinic acid, which along with ferric chloride provided iron in the form of ferric quinate, and succinic acid which are the major sources of growth for the bacteria. Even though magnetosome research has shown promising advances, one of major limitations in its commercialization is its difficulty in cultivation under laboratory conditions.

## I. Introduction

Magnetotactic bacteria (MTB) are widespread, motile, diverse prokaryotes that possess a unique organelle called the magnetosome. Magnetosomes consist of a nano-sized crystal of a magnetic iron mineral that is enveloped by a lipid bilayer membrane [2,7]. In cells of almost all MTB, magnetosomes are organized as a well-ordered chain [20]. The magnetosome chain causes the cell to behave like a motile, miniature compass needle where the cell aligns and swims parallel to magnetic field lines [6]. Magnetosomes consist of magnetic mineral crystals, either magnetite (Fe_3_O_4_) or greigite (Fe_3_S_4_), enveloped by a bilayer membrane composed mostly of phospholipids, called the magnetosome membrane, that contains several proteins not present in the cytoplasmic and outer membranes (OMs) and is unique to MTB [25,27]. Although magnetosome magnetite and greigite crystals can have different morphologies, mature crystals of both minerals generally lie within the single magnetic-domain size range, about 35 to 120 nm, in which they have the highest possible magnetic moment per unit volume [22]. Magnetosomes are usually arranged as a chain within the cell, thereby maximizing the magnetic dipole moment of the cell and causing the cell to passively align along magnetic field lines as it swims [9,18].

These ubiquitous microorganisms represent a morphologically, phylogenetically, and physiologically diverse group of gram-negative bacteria that biomineralize unique organelles called magnetosomes, which are responsible for the cells’ magnetotactic behavior, which is referred to as magnetotaxis [1,19,20]. The origin of magnetotaxis appears to be monophyletic; that is, it developed in a common ancestor to all MTB, although horizontal gene transfer(HGT) of magnetosome genes also appears to play a role in their distribution [10,26]. How the genes involved in magnetotaxis common to all MTB originated and were transferred during evolution is still a matter of debate, although there is evidence that magnetotaxis originated only once, regardless of the composition of the magnetosome crystal, and was then transferred by descent to all groups containing MTB and through Horizontal gene transfer between closely related bacteria [19]. Magnetotactic bacteria thrive in sediments or chemically stratified water columns, where they occur predominantly at the oxic-anoxic interface (OAI), the anoxic regions of the habitat, or both [11]. Although the detection of MTB in samples collected from natural environments is relatively simple to do, MTB is a fastidious group of prokaryotes, and special culture conditions are necessary for their isolation and cultivation.

Nano-magnetite has potential applications in biotechnological and medical fields. These nano-magnetites can be synthesized artificially in laboratories, but, they lack uniform shape and size which could lead to different interaction with different compounds [21]. Since the magnetotactic bacteria are present in stringent localities of the planet, it is difficult to collect samples containing magnetotactic bacteria. Only a handful amount of research and development has occurred in this field. There is no defined technique and media composition for the isolation of these bacteria.

To resolve these issues, we use a natural method where MTB, which naturally synthesize these nano-magnetites through biomineralization are cultivated under laboratory conditions to extract the nano-magnetite [30]. These natural nano-magnetites have uniform shape and size rendering better efficiency in comparison with synthetic nano-magnetites. This method of isolating nano-magnetite from MTB is economical and efficient consisting of high levels of purity and crystallinity. They are thermally stable at physiological temperature and possess high biocompatibility and less toxicity in comparison with synthetic nano-magnetites.

## II. Materials and Methods

### 2.1 Collection and detection of magnetotactic bacteria

#### 2.1.1 Sampling

The MTB are predominantly found in fresh-water, marine, brackish, hyper-saline habitats and sediment regions, depending on the presence of the OATZ [24,25]. In this research, sampling for MTB is based on the collection of the sediment layer that includes and surrounds the OAI and the oxygen transition zones. The soil samples were collected at different depths (0.3, 0.6, 0.9, 1.2, 1.5 m) at different locations (Vijaya Steel Industries – Peenya, Vinayaka Steel centre - Bannergatta Road and Shree Banashankari constructions - Kengeri) by subjecting them to external magnetic fields. The locations were chosen as they were biogeographically suitable for the growth of the bacteria. The magnetic field was imparted by stationing a bar-magnet for an hour in the locality where the sample was about to be collected. Depending on the sample type (e.g., freshwater versus marine habitats), MTB can last from weeks to years, even without the addition of nutrients. In several studies, successions of different magnetotactic bacterial morpho-types have been observed during the enrichment process.

Samples were stored in presence of magnetic influence under dim light at room temperature (25 °C) to avoid proliferation of phototrophic organisms that often leads to a significant decrease or elimination of MTB.

#### 2.1.2 Media Formation

For chemo-organo-heterotrophic growth, the most effective choices appear to be organic acids (e.g., succinate and acetate) and some amino acids, as no MTB have been shown to utilize any other type of organic compound (e.g., carbohydrates) as a carbon source [6]. Iron is required for magnetosome synthesis, and therefore, it must be present in the growth medium. The type of iron source is not critical, however, if it is kept soluble at neutral pH by the presence of either chelating agents particularly if the iron is supplied as Fe(III) or reducing agents that reduce Fe(III) to the much more soluble Fe(II) form [12]. An optimum media was formulated as shown as in Table 1. The vitamin and mineral solutions were formulated based on the requirements for the bacterial growth. Coffee bean solution was used as an alternative for quinic acid and succinic acid. It was prepared by taking 5 – 7 raw coffee beans that were ground in mortar and pestle and then mixed with phosphate buffer. The solution was centrifuged at 2000 RPM for 3 minutes at 4°C. FeCl_3_ was added to the supernatant till a concentrated solution was obtained. This solution was prepared to provide a high concentration of iron in the form of ferric quinate and succinic acid for the growth. To determine relationship between the impact on growth of bacteria and the iron required for its sustained growth, the media was split into Control group with media containing all the necessary media solutions except coffee bean extract and a Test group with all the media solutions including coffee bean extract.

**Table 01.**
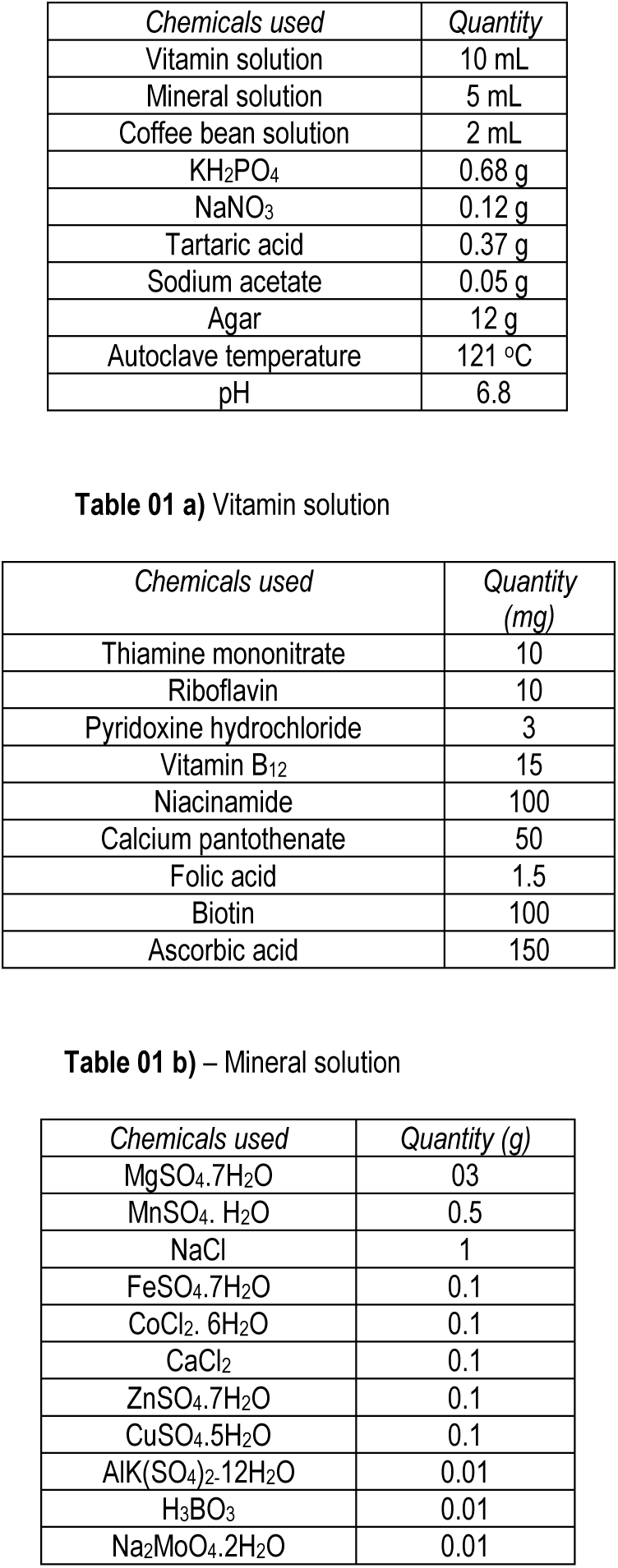
Media composition

The serial-dilution technique was employed as the primary step to analyse growth of the bacteria in the laboratory conditions. The serial dilution was carried out for every depth (0.3, 0.6, 0.9, 1.2, 1.5 m) of the soil sample collected at all the locations. This technique was to confirm the presence of the bacteria and to find the optimum dilution required for the inoculation and cultivation of the bacteria.

### 2.2 Primary detection of MTB

Gram-staining technique was primarily employed to confirm the gram-negative nature of MTB. It was performed for all the dilutions (10^−1^-10^−5^) and at all depths (0.3-1.5 m). Dilutions greater than 10^−3^ did not show satisfactory results in all the depths. Convincing results were obtained at the soil sample collected from 0.6-0.9 m depth with a dilution of 10^−2^. Bar magnets were flanked on either side of the slide and were observed under the microscope. The results showed aggregates on the corners, near the magnetic fields. These aggregates show that the magnetic particles which were present resulted in clumping due to intermolecular attraction and magnetic field.

### 2.3 Cultivation of MTB

#### 2.3.1 Stab-culture

Agar media (50 °C) was mixed with serial diluted bacterial sample and poured into petri dishes and were left to solidify. This was done to provide limited oxygen condition for the media. Using a sterile straight wire, pick a single colony from a freshly streaked plate or from a fresh culture, and stab it deep down into the soft agar several times. Incubate the vial at 37°C for 8–12 h leaving the cap slightly loose. The vial was tightly and store in the dark, preferably at 4°C. The petri-dishes were hermetically sealed and were incubated at 37°C for 24-48 h. This was conducted to analyse the aerobic/anaerobic nature of the bacteria and the concentration of the oxygen required.

#### 2.3.2 Pour-plate method

The media prepared was poured on a petri-plate and the dilution used was 10^−2^ obtained from a depth of 0.6 m, under LAF conditions. Pour-plate was carried on two media groups, one with control group, without coffee bean extract and the other as test group with media having coffee bean extract. The plates were hermetically sealed with paraffin and stored in dark at room temperature for 3-4 weeks under the influence of magnetic field on one side of the petri-dish as shown in the Figure 03. The magnetic field was imparted by placing the disc magnets on the corner of the metal wall. The observation revealed the growth of bacteria only near the zones of magnetic field in test sample. The region of the petri-plate which was not under the influence of magnetic field did not show successful growth of the bacteria. This confirmed the MTB were present in the soil samples and were able to grow successfully in media prepared in the laboratory.

**Figure 01.**
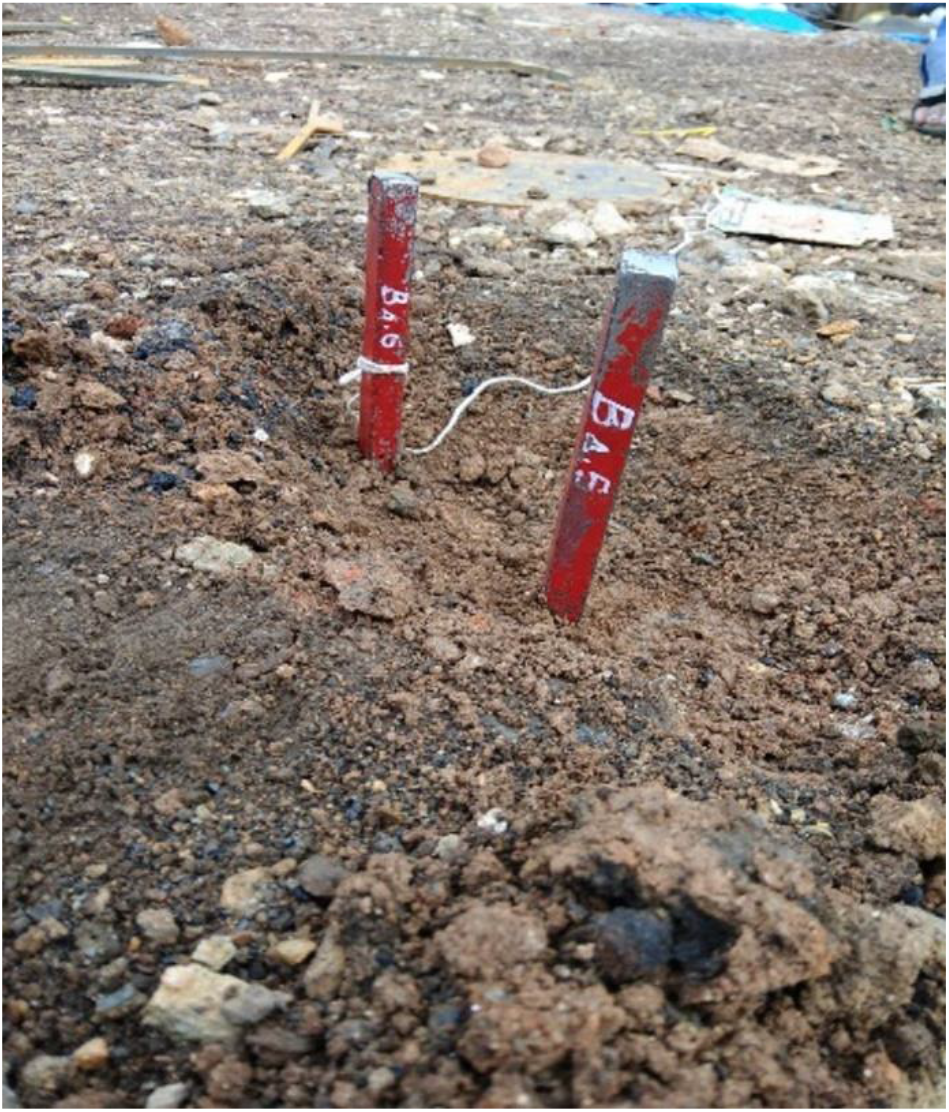
Samples collected iron rich regions.

**Figure 02.**
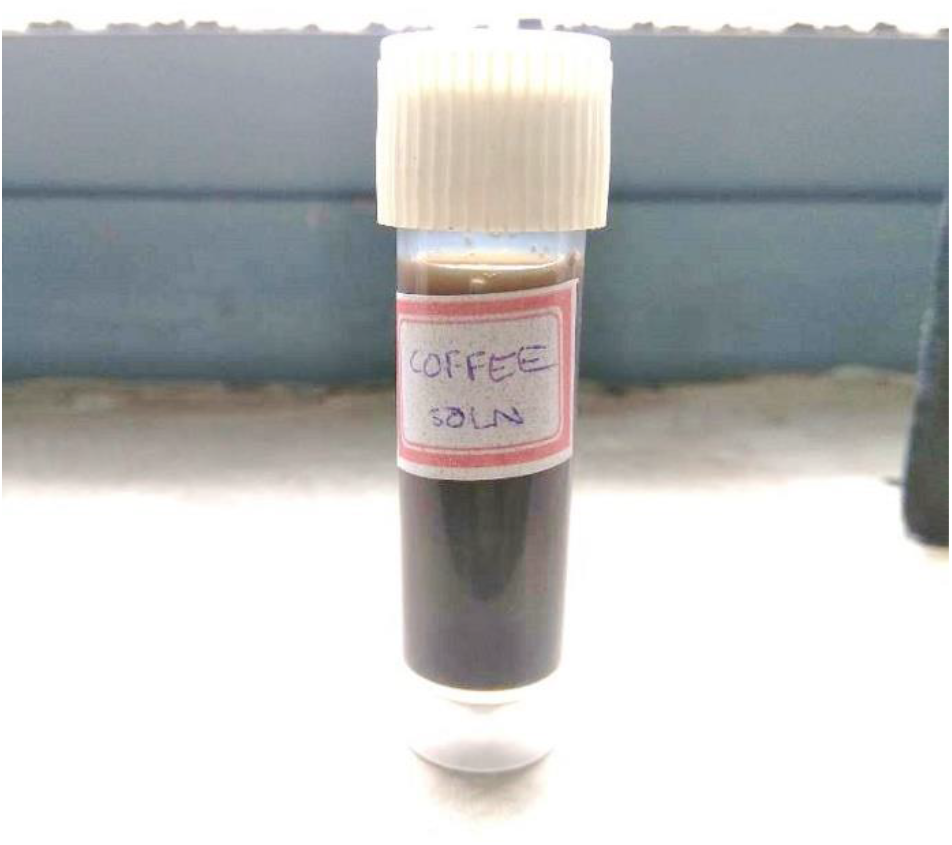
Coffee bean solution

**Figure 03.**
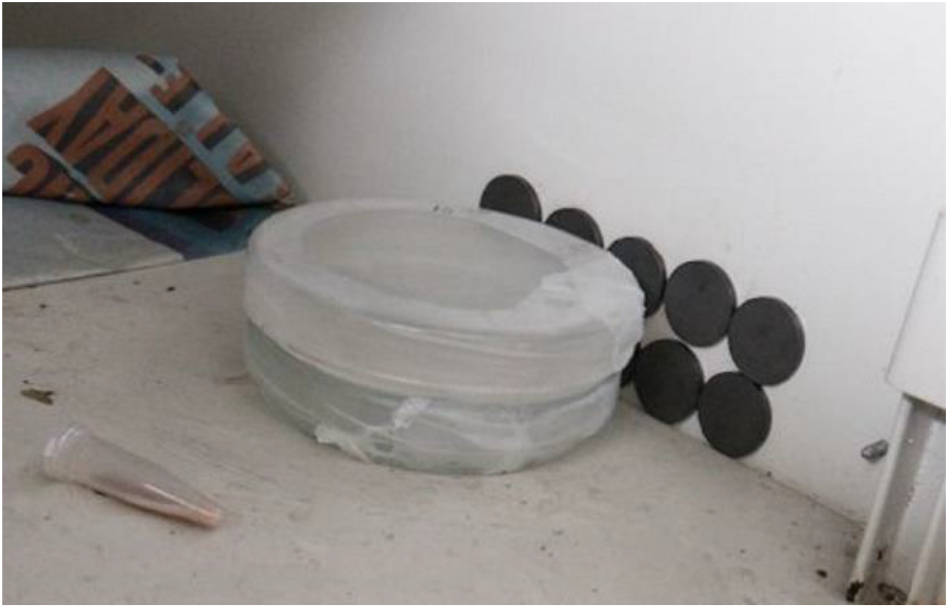
Pour Plate method for control and sample

#### 2.3.3 SEM Analysis

The SEM was used to characterize the bacteria that were obtained from pour-plate method [16]. The parameters from SEM at 5kV, x1k Magnification, working distance = 16.8mm were set to observe the bacteria. The parameters from SEM were set at 6kV, x7.5k Magnification, working distance = 17.2mm to observe the magnetosome chain. The images showed the presence of magnetosome chain and confirmed the bacteria obtained were MTB. The scale taken for the analysis was 50 μm.

## III. Results and Discussion

### 3.1 Results

We observed the presence of bacterial aggregates during gram staining which confirms the magnetic nature of the bacteria. Clumping was observed in the periphery of the petri-dishes due to the influence of magnetic field, as shown in Figure 4 (a). Due to inadequate supply of oxygen, the stab culture showed no growth of bacteria.

**Figure 04.**
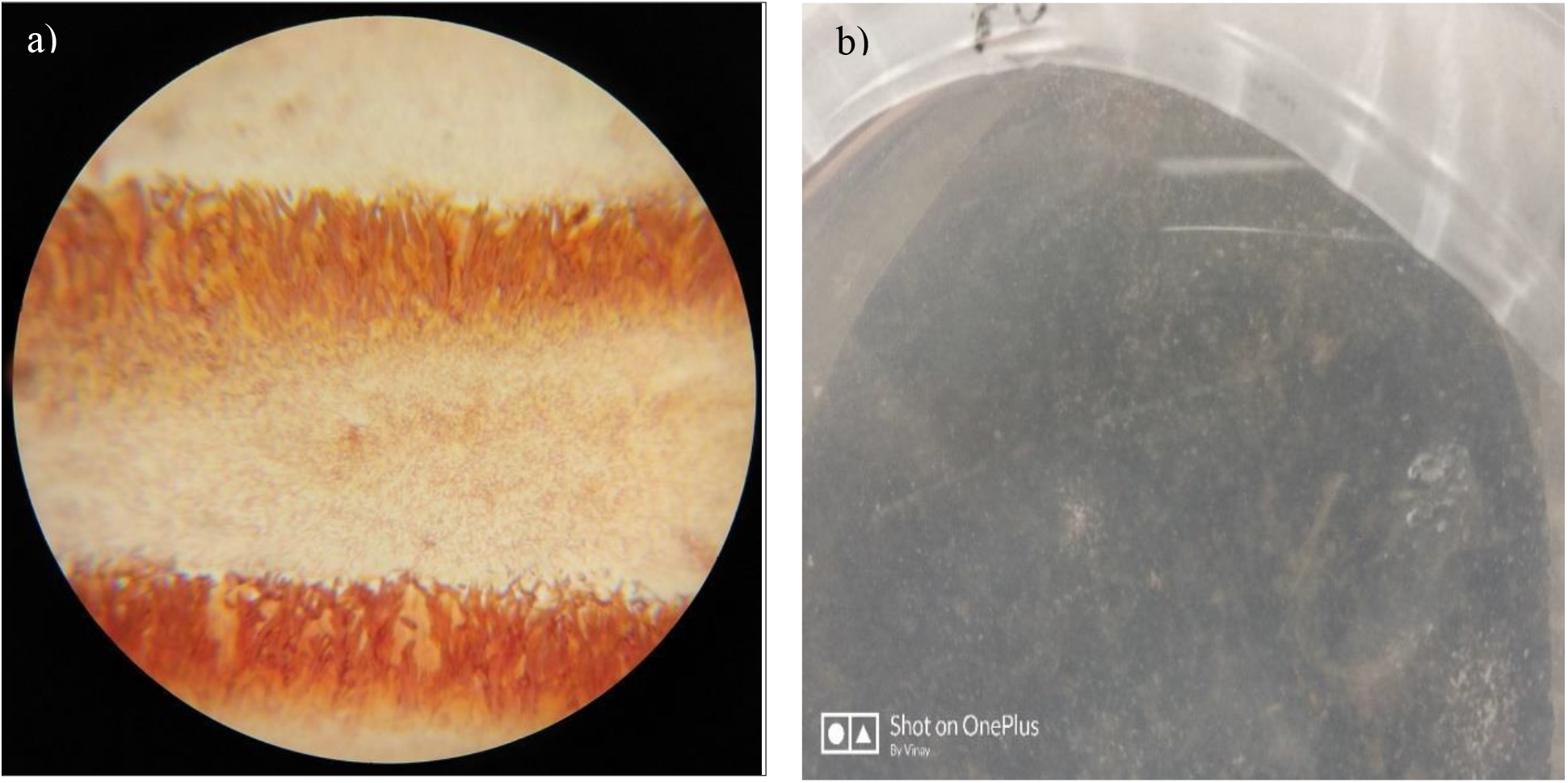
a) Visualisation of gram-staining in the microscope b) Pour-plate in a petri-dish

The pour-plate with the control showed negligible growth of bacteria. The plate with test-sample showed the growth of MTB only in the proximity of magnetic field, as shown in Figure 04 (b). This method confirmed that the bacteria strive and grow only if sufficient iron which was provided in the test media. The coffee bean extract was prepared as an alternative for the chemically produced ferric quinate and to promote use of eco-friendly resources in the growth and development of MTB.

SEM (Vacuum – 6 kV, Magnification – x7.5k, WD – 17.2 mm) analysis confirmed the presence of magnetosome chains in MTB. They also confirmed the shape (rod) of the bacteria and its locomotion by the magnetosome chain [19]. The thin dark line in the images, as shown in Figure 05 (a) and 05 (b), showed the presence of magnetosome chain and confirmed the bacteria obtained were MTB. These chains are produced due to the protein and lipid binding agents and weak intermolecular attraction between the magnetosomes.

**Figure 05.**
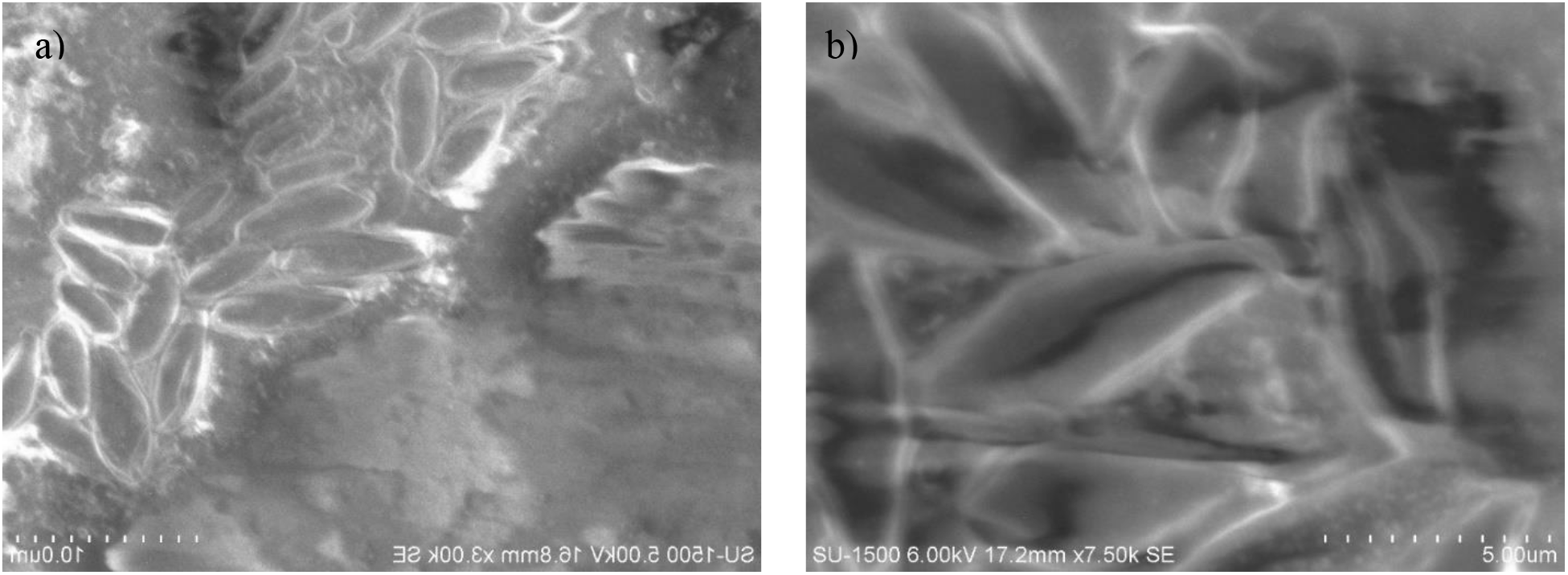
a) Results from SEM at 5kV, x1k Magnification, Working Distance = 16.8mm b) Results from SEM at 6kV, x7.5k Magnification, Working Distance = 17.2mm

### 3.2 Discussion

As the need for iron and its ores (magnetite or gregite) are increased day-by-day in all the fields of development, majorly in fields of biotechnology, nano-magnetites and nano-gregites for various medical, biomedical designs and biproducts [5,33]. MTB, which possess these magnetic chains, are also used in medical field as an alternative for chemotherapies in tumour treatment, carriers for receptors, antigens and antibody, drug delivery [3,4,15,17]. The use of bacteria and nano-magnetite synthesised from natural resources provide comparatively less toxic and chemical reactions when used in medicine [8,23,35]. To use MTB in various applications in future, the need for development of sustainable media and environmental conditions for laboratory growth of MTB is necessary [31,34]. This project represents the successful growth of bacteria from soil and its growth under laboratory conditions with economically and naturally available resources. As the future of biotechnology depends on sustainable development of health and environment, the use of magetotactic bacteria will provide success in future.

The unresolved issues in the isolation of nano-magnetic from MTB where the mechanism of biomineralization to produce nano-magnetite is relatively unknown [32]. Various models have been proposed in this field which are not definitive enough to be applied. Several efforts were carried out to incorporate the genes from MTB in *E. coli*, but the growth and the amount of magnetosome formation were low [22]. The complete genome sequence of MTB is unavailable which a limitation to carry out research on MTB is. Even though magnetosome research has shown promising advances, there are many limitations in its batch cultivation.

Some limitations of MTB are the toxicity of magnetosomes which is supposed to be due to the physical properties of nanoparticles, which have been shown to induce embolism, blockage, and deposition in the body nano-scale size leads to deposition and aggregation of nanoparticles in the body; impurities particularly proteins, nucleic acids, and polysaccharides associated with magnetosomes extracted from bacterial cells, may result in as well membrane proteins of magnetosomes [29]. Moreover, the administration of whole inactive magnetotactic bacteria in humans is not recommended due to the presence of potentially toxic biologic materials such as bacterial DNA or endotoxins. A sample of shortened magnetosomes (obtained by sonication) behaves magnetically in a different manner, showing that both the energy loss and the specific absorption rate are reduced, and thereby indicates a variation in the heating process [14].

Even though magnetosome research has shown promising advances, the only limitation in its commercialization is the availability of magnetosomes. In consideration of its gravity, the cells in microgravity do not show magnetotaxis and discernible chains of magnetosomes suggest that in the terrestrial environment, MTB use magnetosomes to enhance their response to gravity. Therefore, only a limited number of MTB can be isolated as a pure culture, hence due to the low cell densities, the sufficient numbers of magnetosomes cannot be purified which further hampers industrial applications. The emerging opportunities relating to MTB are high-data storage, MRI, treating disease by a mannose-binding lectin, DNA discrimination analysis, drug delivery, they act as carriers for enzymes, nucleic acids, antibodies, biosensors and waste-water treatment [13,28].

## IV. Acknowledgement

This project was supported by Dr. Ashwani Sharma, Assistant Professor, Dr. Nagashree N Rao, Associate Professor, Department of Biotechnology, RVCE, who provided insight and expertise that greatly assisted the project.

We wish to express our sincere gratitude to Dr. Bhuvaneshwara Babu T, Professor, and HOD of Physics Department, RVCE for providing us with bar magnets, which helped us immensely in the collection of the soil sample.

We sincerely thank Dr. Satheesh Babu Gandla, Interdisciplinary research center, RVCE and Dr. Ravishankar H N, Associate Professor, Department of Biotechnology, RVCE for their guidance and encouragement in carrying out the project work.

We also thank Mrs. Puneetha T R, Lab Instructor, Department of Biotechnology, RVCE and Ms. Aishwarya Bhat, Student, Department of Biotechnology, RVCE for providing providential support during the project.

We would like to include a special note of thanks for Dr. Pushpa Agrawal, HOD, Department of Biotechnology, RVCE, for providing us with sufficient facilities required for the completion of the project.

We express our sincere gratitude to the Principal of RVCE, Dr. K N Subramanya for providing us with the opportunity to carry out our project.

We are grateful to our parents for supporting us in all our endeavors.

